# Comparative analysis of the modulation of perineuronal nets in the prefrontal cortex of rats during protracted withdrawal from cocaine, heroin and sucrose self-administration

**DOI:** 10.1101/2020.04.07.029868

**Authors:** David Roura-Martínez, Paula Díaz-Bejarano, Marcos Ucha, Emilio Ambrosio, Alejandro Higuera-Matas

## Abstract

Relapse into drug use is a significant problem for people recovering from addiction. The ability that conditioned cues have to reinstate and reinvigorate drug-seeking is potentiated over time (incubation of seeking), posing an additional difficulty for maintaining abstinence. While the prefrontal cortex has been involved in the incubation phenomenon and the extracellular matrix, perineuronal nets (PNN) in particular, may play a vital role in brain plasticity associated to drug relapse, there are no comparative analyses between different drug classes and natural reinforcers. Here, we compare the effects of early (1 day) and protracted (30 days) withdrawal from to cocaine, heroin and sucrose self-administration on the PNN content of different territories of the prefrontal cortex of male Lewis rats. Our results show that cocaine self-administration and protracted withdrawal decreased PNN content in the prelimbic cortex. Also, heroin self-administration increased PNN content in the infralimbic cortex, but this effect was lost after 30 days of withdrawal. Heroin self-administration also decreased PNNs in the insula, an effect that remained even after protracted withdrawal from the drug. Finally, the self-administration of sucrose-sweetened water decreased PNN content in the dorsomedial prefrontal cortex and increased PNNs in the insular cortex, which was still evident after protracted withdrawal. Our results show that three different rewards with specific pharmacological and physiological actions differentially modulate PNNs in specific areas of the rodent prefrontal cortex with potential implications for the incubation of seeking phenomenon.

## INTRODUCTION

Relapse is a significant problem in the treatment of addiction (Bossert et al., 2005). Multiple factors contribute to relapse in humans and in animal models, including the exposure to cues and environments that have been consistently paired with the effects of drugs of abuse in the past (O’Brien, 2008). The ability of conditioned cues to elicit drug craving is potentiated (incubated) over time, a phenomenon observed in humans (Bedi et al., 2011; P. Li et al., 2015; Parvaz et al., 2016; Wang et al., 2013) and animas, using a wide variety of drugs and other reinforcers (Abdolahi et al., 2010; Aoyama et al., 2014; Bienkowski et al., 2004; Blackwood et al., 2018; Grimm et al., 2002, 2001; Kirschmann et al., 2017; Krasnova et al., 2014; Shalev et al., 2001; Shepard et al., 2004).

During the last decade, several laboratories have suggested that the extracellular matrix (ECM), and in particular perineuronal nets (PNN) play a vital role in brain plasticity, with an impact on learning and memory processes, as well as in different psychopathologies, including addiction (Lasek et al., 2018; Sorg et al., 2016). PNNs are specialized aggregations surrounding the cell bodies and proximal neurites of specific neurons, mostly parvalbumin GABAergic interneurons, that stabilize synapses in the adult brain. It has been shown that the exposure to (either passively or self-administered) and abstinence from drugs of abuse alters the structure of PNN in various brain regions and, conversely, the manipulation of PNNs affects drug self-administration and extinction learning (Lasek et al., 2018).

The prefrontal cortex (PFC) is involved in certain aspects of addiction (Everitt and Robbins, 2016; George and Koob, 2010) and the PNNs located in this region play a role in addictive disorders. For example, continued ethanol consumption (passive administration) or a hypercaloric diet that leads to learning and memory deficits, induce alterations in the PNNs of the PFC (Coleman et al., 2014; Reichelt et al., 2019). Also, the pharmacological manipulation of the ECM in the PFC affects both locomotor sensitization and the conditioned place preference induced by psychostimulants (Mizoguchi et al., 2007; Slaker et al., 2015). Although the PFC also has been suggested to play a role in the incubation phenomenon (Shin et al., 2018), it is still unknown whether the contingent self-administration of drugs or natural rewards and the subsequent protracted abstinence from them, alters PNNs in the different regions of the PFC. To the best of our knowledge, only two studies, Slaker et al. (2016) with sucrose, and Van Den Oever et al. (2010) with heroin, have examined this possibility. However, some limitations in the design of these experiments difficulted the conclusions that can be drawn from them. More specifically, in the study by Slaker et al., the rats underwent a relapse/extinction test before the analyses, which could have affected the composition of the ECM by itself. On the other hand, in the Van Den Oever report, the components of the ECM were only examined after three weeks of withdrawal (but not acutely, i.e. after 24 hours), and only in the medial PFC.

In considering all these pieces of evidence, the main goal of this work was to study the modulation of PNNs in different subregions of the PFC, using three rewarding substances with different pharmacological effects – cocaine, heroin and sucrose – at two different abstinence points (1 and 30 days) after a contingent consumption protocol (self-administration) that is known to induce incubation of seeking.

## MATERIALS AND METHODS

### Animals

Tissue for PNNs analyses was derived from 84 Lewis male rats purchased from Harlan International Ibérica weighing 300-320g at the beginning of the experiments (6 samples were lost due to storage problems). Upon arrival, the animals were housed in groups of three in the vivarium at a constant temperature (20 ± 2 °C) and on a 12h:12h light:dark cycle (lights on at 08:00 a.m.), with food (Panlab, commercial diet for rodents A04/A03) and water available ad libitum. Animals were maintained and handled in accordance with European Union Laboratory Animal Care Standards (2010/63/EU). In the spirit of reducing the number of animals used in the experiments, these rats belonged to a subset of animals used in a study previously published by us, focused on the incubation of seeking of cocaine, heroin or sucrose and examining the glutamatergic, GABAergic and endocannabinoid systems (Roura-Martínez et al., 2019). Of note, these animals showed a self-administration pattern that was indistinguishable from the behaviour of the rats that underwent extinction testing and that served us to verify that we would indeed observe the incubation phenomenon under our conditions. Detailed methodological information and behavioural results can be found in this previous work (Roura-Martínez et al., 2019).

### Experimental design

Two batches of rats were used, one submitted to jugular catheter surgery (for cocaine, heroin or saline self-administration) and the other left intact (for sucrose or water self-administration). After self-administration sessions, the rats underwent 1 or 30 days of forced withdrawal (with regular handling) without extinction testing. The animals that received intravenous administration were segregated into six groups (n=8 rats per group, 3 substances -heroin, cocaine, saline- x 2 withdrawal periods), while the rats that consumed the substance orally (sucrose-sweetened water, or tap water) were distributed in four groups (n=9 rats per group, 2 substances x 2 withdrawal periods).

### Surgical procedures

An intravenous (i.v.) polyvinylchloride tubing (0.064 mm i.d.) catheter was implanted into the right jugular vein at approximately the level of the atrium and passed subcutaneously to exit the midscapular region. Surgical procedures were performed under isoflurane gas anaesthesia (5% for induction and 2% for maintenance) and buprenorphine analgesia. After surgery, the rats were allowed to recover for 7 days and a nonsteroidal anti-inflammatory drug (meloxicam, Metacam™: 15 drops of a 1.5 g/mL solution per 500 mL of water) was added to the drinking water. The catheters were flushed daily with 0.5 mL of gentamicin (40 mg/mL) dissolved in heparinized (100 IU/ml) saline in order to prevent infection and to maintain patency.

### Self-administration

All the self-administration sessions were performed in Skinner boxes (Coulbourn Instruments or Med-Associates) and they were monitored with Med-PC software. The house light was off during the sessions, although we allowed some environmental light from the room (the door of the sound-attenuating cubicle was left ajar) in order to keep the light:dark cycle. Two levers were used, an active (L1) and inactive lever (L2). Each time L1 was pressed by the animal (LP1; fixed-ratio 1), a pump outside the box was switched on for 5 seconds and either 100 μL of the drug or saline solution was infused through the catheter, or 100 μL of sucrose solution or water was dispensed into a receptacle placed in between the levers. A cue-light over the active lever also switched on for 10 seconds at the same time. Subsequently, there was a time-out period of 40 seconds in which there were no programmed consequences, although the responses at each lever were recorded. Cocaine, heroin or saline self-administration sessions lasted 6 h each day. Rats orally self-administering sucrose or water were subjected to shorter sessions (2h/day). The doses per injection used in the experiments were 0.075 mg/kg of heroin; 0.75 mg/kg of cocaine-HCl; and 10% w/v sucrose (Sigma-Aldrich S1888), diluted in 0.1 mL of saline (0.9% NaCl physiological saline Vitulia-ERN, for the intravenous infusions) or tap water. In the first two self-administration sessions, two sucrose pellets were placed on the active lever to facilitate the acquisition of self-administration behavior.

### Animal sacrifice

One or 30 days after the last self-administration session, the animals were weighed and sacrificed by decapitation between 11:00 and 13:00 a.m. The brain of the rats was quickly extracted and submerged in isopentane chilled on dry ice for ten seconds and to be then stored at −70 °C.

### Dissection

Each brain was embedded in TissueTek (Sakura, 4583) and kept at −20 °C in a cryostat chamber (Microm, Cryostat HM 500O). After one hour, 40μm coronal slices of the PFC (+3.7mm from Bregma) were obtained and placed on adhesive slides (Thermo Scientific, SuperFrost® Plus, Menzel Gläser). After air-drying, the slides were stored at −30°C until further use.

### Fluorescent staining of perineuronal nets

The samples were thawed at 4°C for 30min and then kept at room temperature (RT) for other 30min, then incubated in cold (−20°C) post-fixation solution (4% w/v formaldehyde in PBS 0.1M pH 7.4) for 15min. They were allowed to air dry at RT for 20min and washed 3×15min in PB. The samples were then incubated overnight with WFA-FITC (Vector FL1351) 1:250 in PB with 0.25% v/v Triton X-100. The incubation was carried out in a humid chamber in the dark, at 4°C on a flat surface. The next day, three 15-minute washes were performed in PB. Each slice was incubated in 100μL of DAPI (Sigma-Aldrich D9542) 1:1500 in PB at RT in the dark for 5min and then two 15min washes were performed in PB. The slices were covered with ProLong Gold antifade reagent (Thermo Fischer P36930) and coverslips and left to dry 24h in the dark.

### Fluorescence microscopy

Images from different regions of the prefrontal cortex (both hemispheres) were obtained with an Olympus U-RFL-T fluorescence microscope (Olympus DP70 camera) at x10, x20 and x40. Two x40 images were taken per animal that comprised all the layers of the insular (IC), anterior cingulate (ACC), dorsal prelimbic (dPL), ventral prelimbic (vPL) and infralimbic cortices (IL). An additional image per animal was obtained that included the ventral (vOFC) and lateral orbitofrontal cortex (lOFC), according to the atlas of Paxinos and Watson (2007) (see Figure 1).

**Figure 1:**
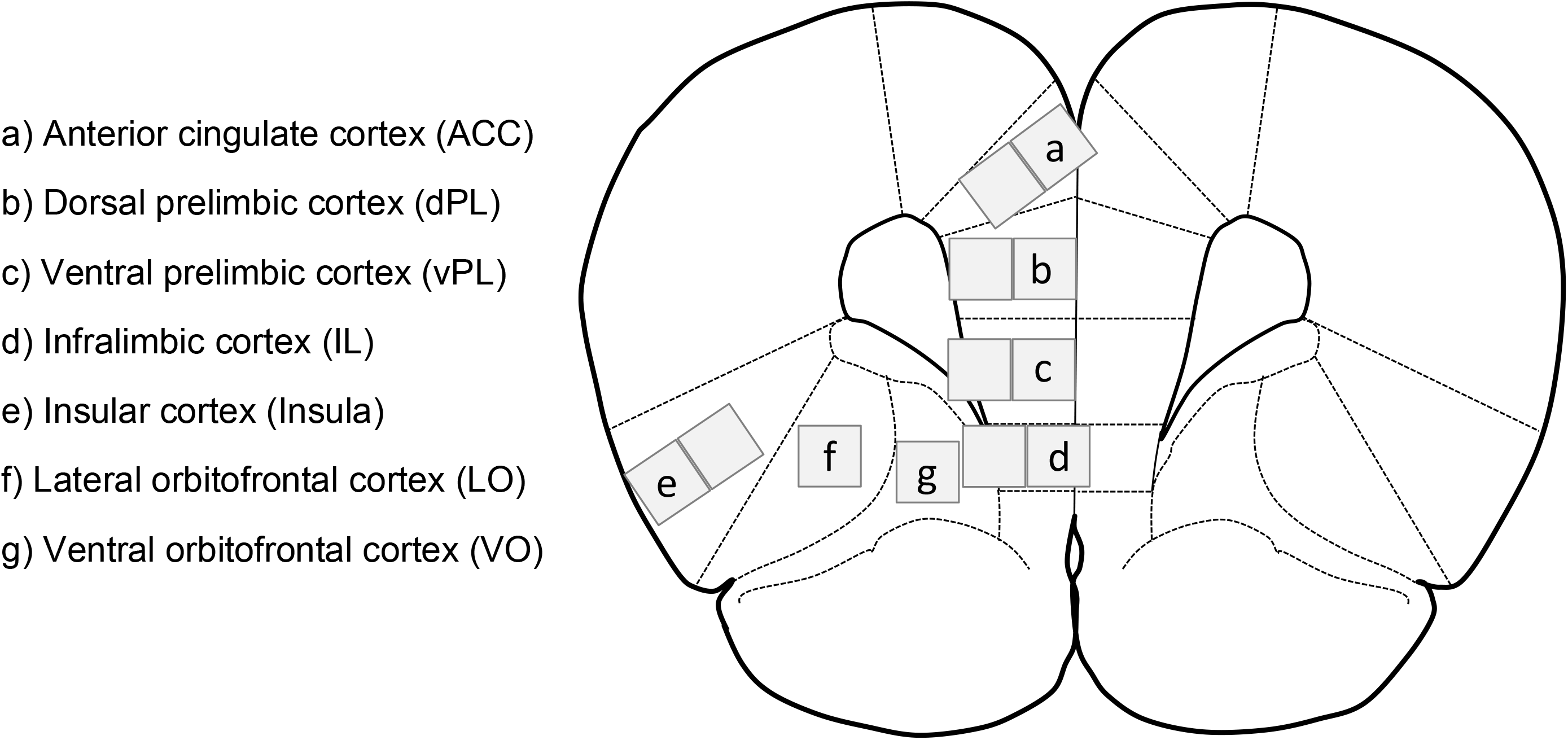
Locations of the micrographs used for the analysis.

### Analysis of PNN

The images were analysed with the free software Fiji (ImageJ). From each region, the number of perineuronal nets per square micrometre (PNN/μm2) was calculated, regardless of their intensity. We first assayed two strategies to deal with the high variability in PNN presence: (1) to normalize to the total number of PNNs of each subject and then to perform the ANOVA (Sidak post-hoc) and (2) to perform an ANCOVA using the total number of PNN of each subject as covariate. To deal with missing data when calculating the total value per subject in our multivariate analyses, we applied an imputation procedure based on bootstrapping (implemented in IBM Statistics). These imputed data were only used for normalization purposes but were not included in the final analyses. The ANCOVA approach had to be discarded because the covariate showed a significant interaction with the treatment and/or the time of abstinence. Thus, PNNs in each area were analysed by factorial ANOVA as the proportion of the total PNNs expressed by each animal in the entire PFC. Given that the imputation procedure generates five different models for the lost values, the results are presented as ranges (mean±SEM). Statistical significance was set at 0.05.

## RESULTS

A detailed analysis of the self-administration behaviour of the rats from which the tissue was obtained can be found in our previous report (Roura-Martínez et al., 2019). Briefly, rats undergoing 1 or 30 days of withdrawal displayed similar self-administration behaviour within each of the rewards tested, with the following values (mean±SD) of cumulative active lever presses (p>.30, unpaired t-test): saline wd1 36.9±17.6, wd30 28.9±13.5; cocaine wd1 166.4±90.5, wd30 180.5±70.3; heroin wd1 545.5±368.5, wd30 668.4±440.6; water wd1 23.0±8.7, wd30 23.6±10.7; sucrose wd1 394.8±141.7, wd30 377.6±129.2.

Table 1 shows the results of the statistical analysis of the main factors manipulated here and their interaction, for all the areas of the PFC that were analysed.

**TABLE 1.**
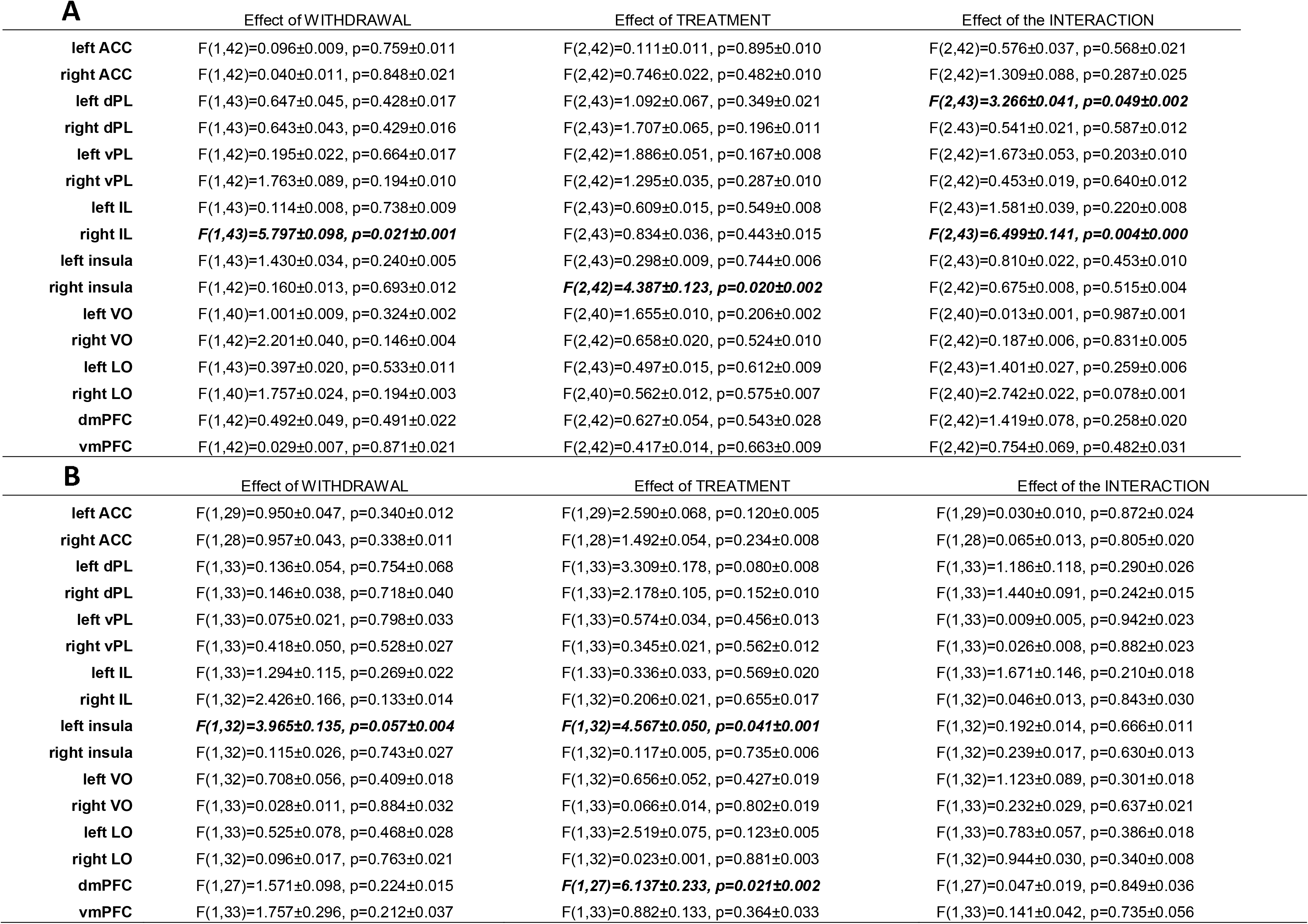
ANOVA of the PNN content in different subregions of the prefrontal cortex in the saline, cocaine and heroin experiment (A) and water/sucrose experiment (B). Data are expressed as the range of the 5 imputation models used to generate the normalized values. In bold, the significant values that are represented in the figures. ACC: anterior cingulate cortex; dPL: dorsal prelimbic cortex; vPL: ventral prelimbic cortex; IL: infralimbic cortex; LO: lateral orbitofrontal cortex; VO: ventral orbitofrontal cortex; dmPFC: dorsomedial prefrontal cortex; vmPFC: ventromedial prefrontal cortex.

### Cocaine self-administration and protracted withdrawal decreased PNN content in the prelimbic cortex

The analysis of the dorsal prelimbic cortex revealed a significant *Substance* x *Withdrawal Day* interaction (*left hemisphere*: F_2,38_=3.266±0.041, p=0.0492±0.0017, η^2^=0.138±0.002). The simple effects analysis revealed a significant decrease in PNN content in the group of rats that consumed cocaine and underwent 30 days of withdrawal as compared to their saline control (*p*=0.0358±0.0026) and to the cocaine group undergoing 1 day of withdrawal (*p*=0.0179±0.0012) (see Figure 2 A,B).

**Figure 2:**
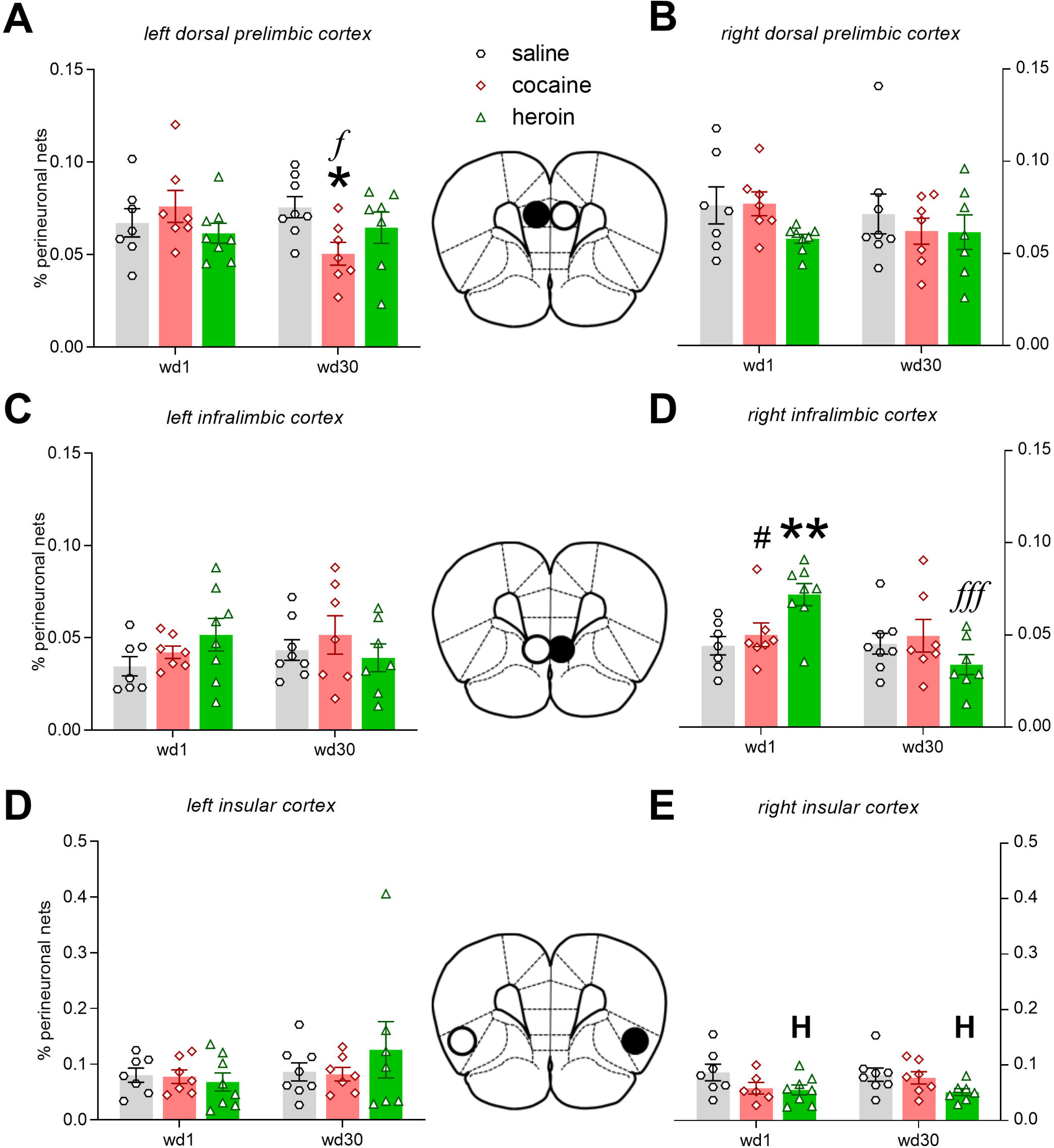
Mean±SEM of the perineuronal net content (expressed as a percentage of the total PNN in the entire PFC of the animal) in animals exposed to saline, cocaine and heroin under a 6h self-administration regime and undergoing 1 or 30 days of forced withdrawal. The black circle in the brain scheme depicts the location of the significant effects. * p<0.05 ** p<0.01 compared to corresponding saline group. *f* p<0.05 *ff* p<0.01 against the corresponding WD1 group. # p<0.05 against the corresponding heroin group. **H** p<0.05 general effect of heroin (found after the post-hoc analysis of the *Substance* effect).

**Figure 3:**
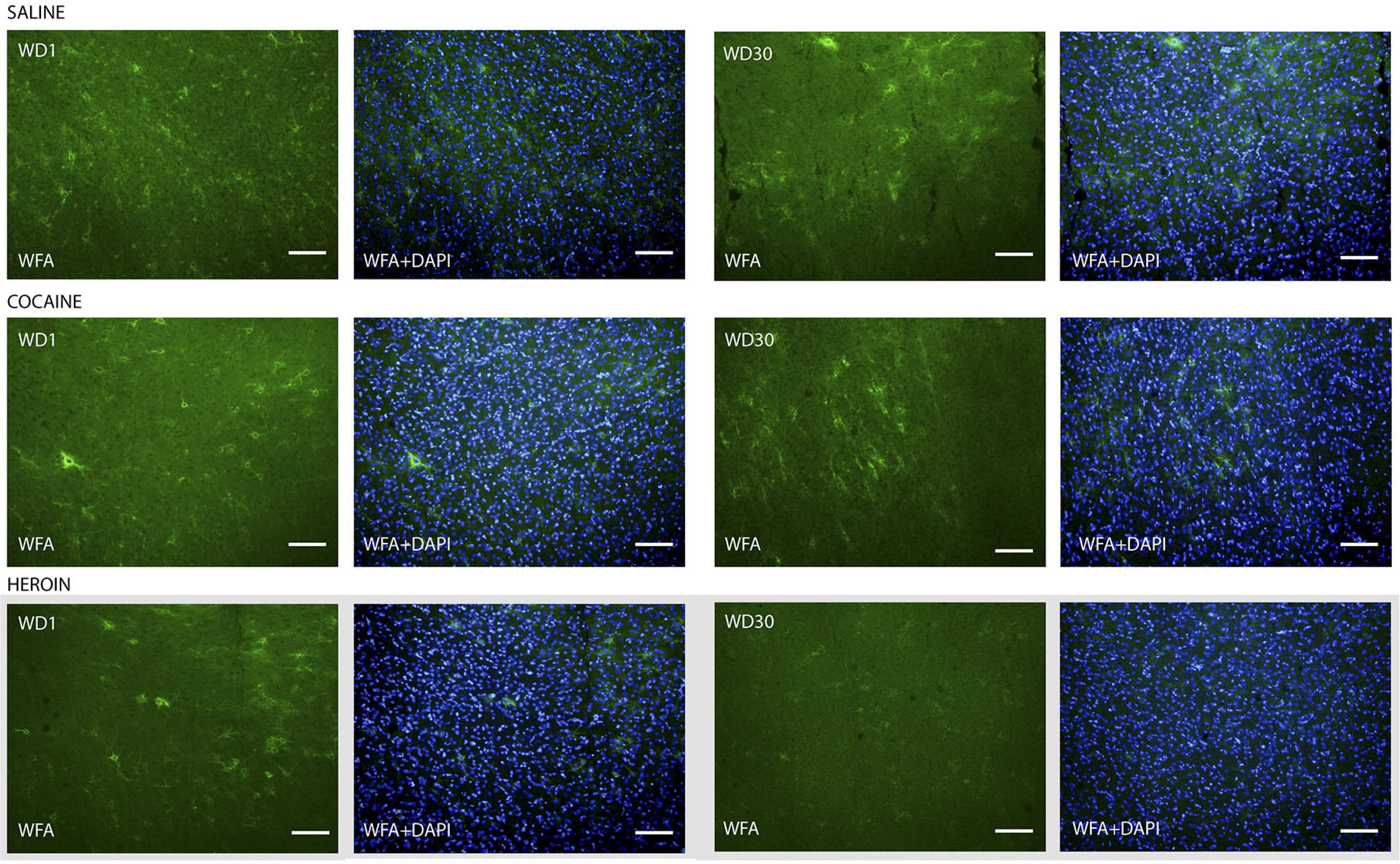
Representative photomicrographs taken in the infralimbic cortex (right hemisphere) of perineuronal nets stained with *Wisteria Floribunda* (WFA) hemagglutinin and the merged image with nuclei stained with DAPI. The grey box indicates the representative images belonging to the groups where the significant effects are found. Scale bar = 50 μm.

### Heroin self-administration temporally increased PNN content in the infralimbic cortex and decreased PNNs in the insula

The analysis of the infralimbic cortex revealed a significant S*ubstance* x *Withdrawal Day* interaction (*right hemisphere*: F_2,38_=6.499±0.141, p=0.0038±0.0004, η^2^=0.219±0.0004). Our simple effects analyses revealed a significant increment in the group of rats that self-administered heroin and underwent 1 day of withdrawal as compared to its saline control (*p*=.0094±.001), the cocaine group of the withdrawal time (p=.0479±.0026), and the heroin group undergoing 30 days of withdrawal (p=0.0001±0.0001). Also, a general effect of the *Withdrawal Day* was observed (F_1,38_=5.797±0.098, p=0.0211±0.00103, η^2^=0.0979±0.0019) (see Figure 2 C, D).

Our analysis of the insular cortex revealed a *Substance* effect (*right hemisphere*: *F*2,37=4.3867±0.1233, *p*=.0199±.00196, η^2^=.1862±.00426). The *post-hoc* analyses revealed a significant decrease after heroin consumption as compared to the saline control group (p=0.0164±0.00173) (see Figure 2 E, F).

### Sucrose consumption decreased PNN content in the dorsomedial prefrontal cortex and increased PNNs in the insular cortex

The analysis of PNN content in the entire dorsomedial prefrontal cortex (i.e. the sum of the values of the anterior cingulate cortex, the dorsal and ventral prelimbic cortices and the infralimbic cortex from both hemispheres) revealed a significant *Substance* effect whereby there was a long-lasting decrease in PNN content after sucrose self-administration (F_1,24_=6.1368±0.23263, p=0.0212±0.00224, η^2^=0.1958±0.00562) (Figure 4 A). However, in the insular cortex, we observed a significant and long-lasting increment in the PNN content after sucrose self-administration (*left hemisphere*: F_1,29_=4.5672±0.04996, p=0.0412±0.00105, η^2^=0.1245±0.00143), as well as a general decrease due to the effect of *Withdrawal Day* (F_1,29_=3.9650±0.13537, p=0.0566±0.00399, η^2^=0.1080±0.0034) (see Figure 4 B, C).

**Figure 4:**
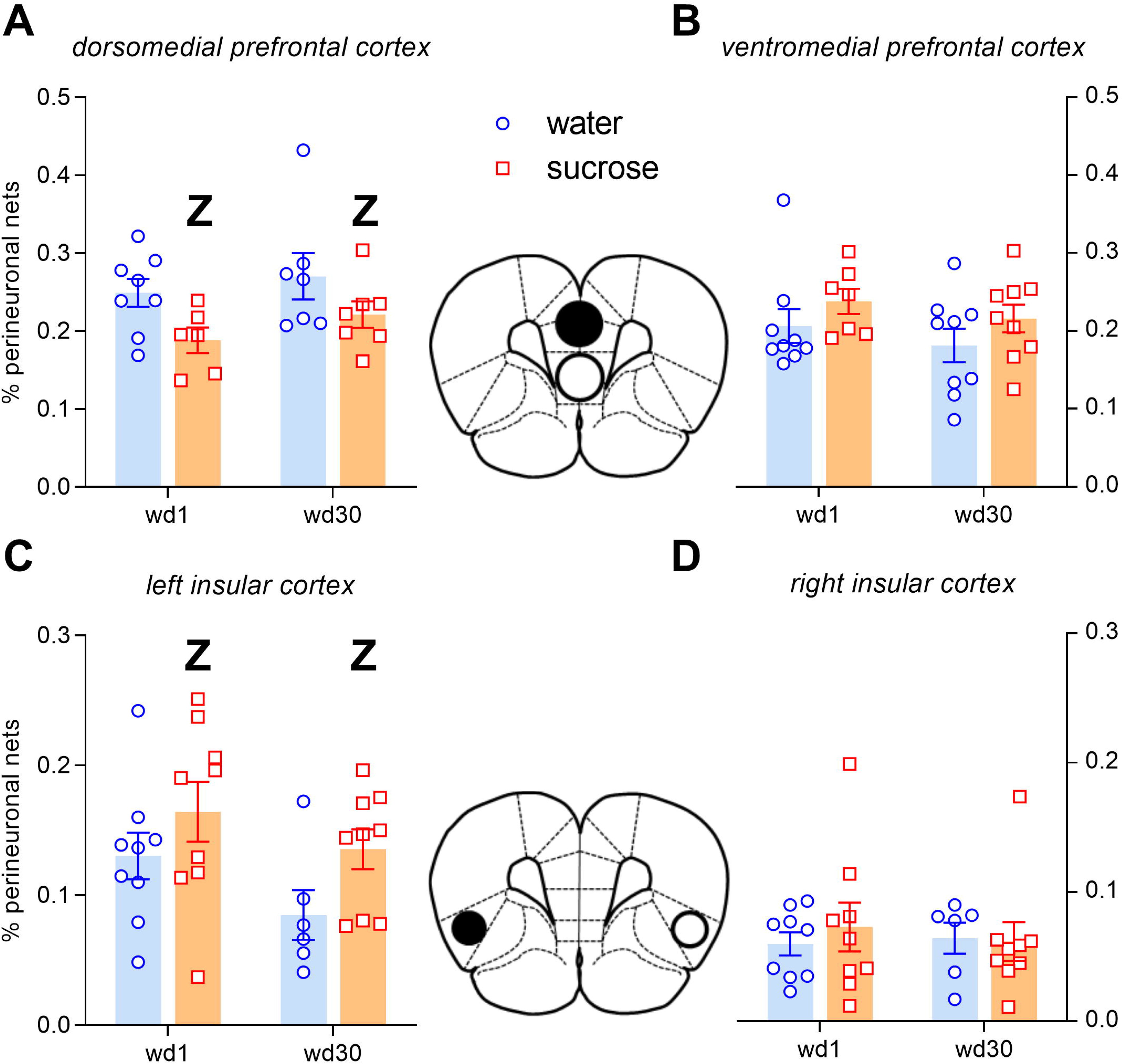
Mean±SEM of the perineuronal net content (expressed as a percentage of the total PNN in the entire PFC of the animal) in animals exposed to water and sucrose-sweetened water a 2h self-administration regime and undergoing 1 or 30 days of forced withdrawal. The black circle in the brain scheme depicts the location of the significant effects. **Z** p<0.05 general effect of sucrose (found after the post-hoc analysis of the *Substance* effect).

**Figure 5:**
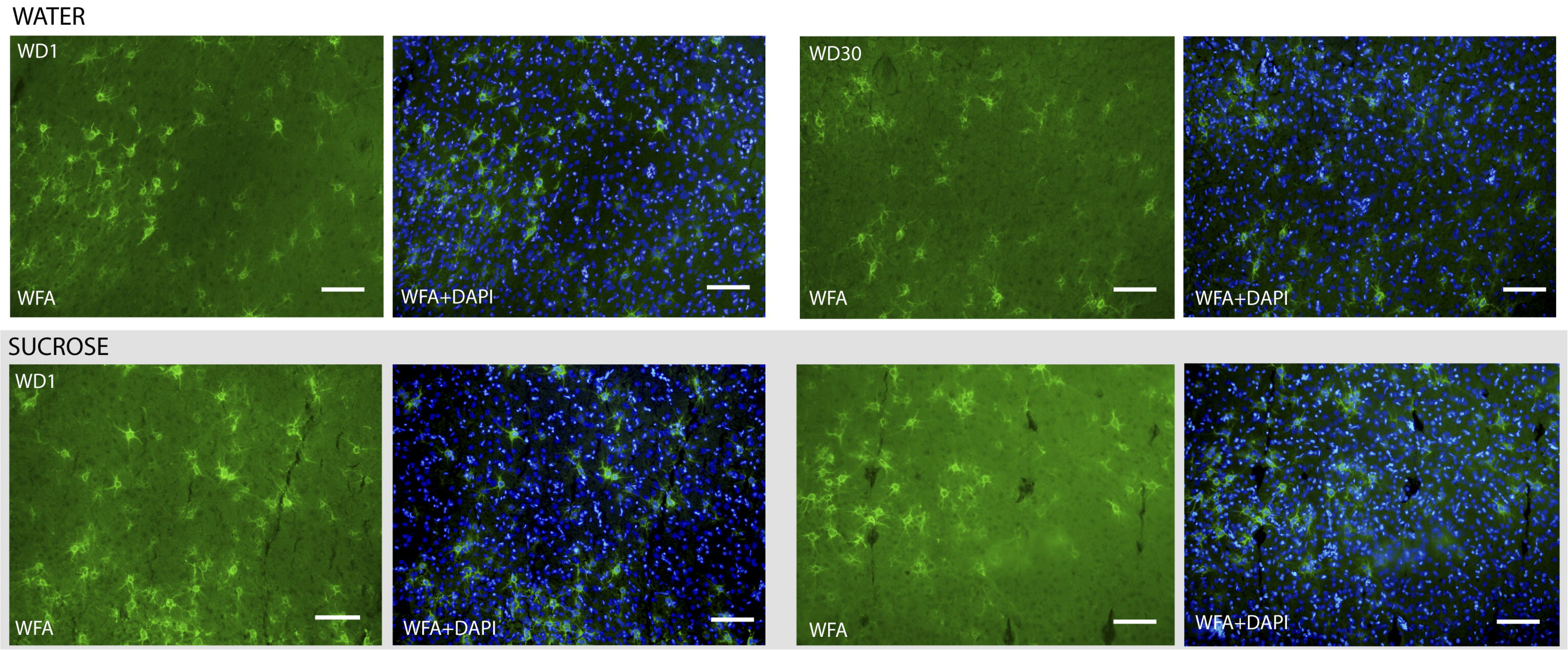
Representative photomicrographs taken in the insular cortex (left hemisphere) of perineuronal nets stained with *Wisteria Floribunda* (WFA) hemagglutinin and the merged image with nuclei stained with DAPI. The grey box indicates the representative images belonging to the groups where the significant effects are found. Scale bar = 50 μm.

## DISCUSSION

In this work, we have performed the first comparative analysis of the effects of early and late withdrawal from three rewarding substances (a natural reward and two drugs of abuse with different pharmacological profiles) in the PNN content of specific subregions of the medial prefrontal cortex of the rat. A general effect of the exposure to these three rewards was a down-regulation on the PNN content. However, there were interesting substance-, region- and hemisphere-specific effects and also precise modulations that depended on the length of withdrawal. For example, the PNN decrease associated with cocaine only occurred in the dorsal prelimbic region and exclusively after protracted withdrawal. At the same time, in the case of heroin, a biphasic modulation was observed in the left infralimbic cortex that did not occur in the right insula where PNNs were decreased at early heroin withdrawal and remained down-regulated after 30 days of abstinence. The only exception was the left insula where sucrose self-administration increased PNN content, and this effect persisted even after 30 days of withdrawal.

### Decreased perineuronal net content in the medial prefrontal cortex

We found a differential pattern of alterations in the content of PNNs in medial PFC, depending on the substance self-administered by the rats. The PNN content decreased in the left dorsal prelimbic cortex after 30 days of withdrawal from cocaine. Also, we observed increased PNN levels after heroin self-administration in the right infralimbic cortex that decreased to control levels after one month of withdrawal from heroin. Finally, there was a decrease in PNN levels in the dorsomedial prefrontal cortex after self-administration of sucrose that was still evident after 30 days of withdrawal.

Previous self-administration experiments showed that the content of PNNs in prelimbic or infralimbic cortices did not change after short (3 days) withdrawal from nicotine (Vazquez-Sanroman et al., 2017). However, another experiment showed that PNNs decreased in the medial prefrontal cortex after protracted (21 days) withdrawal from heroin (Van den Oever et al., 2010). This last result is in line with our cocaine self-administration experiments, where we observed decreases only after one month of withdrawal. Moreover, after heroin self-administration, we also found decreases after 30 days of withdrawal; however, these decrements were only evident in the comparison from a previous increase observed at early withdrawal, a timepoint not measured in the Van Den Oever et al. report. Finally, Slaker et al. (2016) found that the PNN content did not change after short (1 day) or protracted (30 days) withdrawal from sucrose self-administration, which is in contrast to our results since we did find decreases in the dorsomedial prefrontal cortex. Some differences between the studies could explain this discrepancy: 1) Slaker et al. did not measure the total content of perineuronal nets in the dorsomedial prefrontal cortex; and 2) they performed an extinction test before the sacrificing the animals, which may have reversed the decrease in the content of PNNs in the medial prefrontal cortex after protracted abstinence (Van Den Oever et al., 2010). Thus, our results suggest that protracted withdrawal after a self-administration protocol is associated to a decrease in PNN content in the medial prefrontal cortex, in a perdurable (sucrose) or a biphasic (cocaine, heroin) manner.

The implication of the individual prefrontal cortical subregions in the incubation phenomenon is not clear, as highlighted by the contradictory results of the pharmacological manipulations performed in the different subdomains of the medial PFC during the incubation of seeking of psychostimulants (Ben-Shahar et al., 2013; Gould et al., 2015; Koya et al., 2009; X. Li et al., 2015; Miller et al., 2017; Müller Ewald et al., 2018; Shin et al., 2018; Swinford-Jackson et al., 2016; Szumlinski et al., 2018). In spite of the controversy, PNNs in the medial PFC seem to be associated with a decreased seeking response. For example, by pharmacologically interfering with the degradation of the ECM (with an i.c.v. injection of an inhibitor of metalloproteinases), Van Den Oever et al., (2010) showed that the levels of several components of ECM in the medial PFC increased, along with a concomitant decrease in heroin seeking. In addition, another study showed that environmental enrichment increased the levels of PNNs in the prelimbic and infralimbic cortices and decreased sucrose seeking (Slaker et al., 2016).

The decreases in PNN levels after protracted withdrawal that we report here are accompanied by changes in neurotransmitter receptors in the same animals (Roura-Martínez et al., 2019). First, we have observed an increment in the gene expression of NMDA receptor subunits in the dorsomedial prefrontal cortex after cocaine withdrawal, similar to the results reported in the medial prefrontal cortex by Ben-Shahar et al., (2009). We also have documented increases in the gene expression of extrasynaptic subunits of GABA_A_ receptors in the ventromedial prefrontal cortex after heroin withdrawal (Roura-Martínez et al., 2019); and a general depletion in the expression of glutamatergic, GABAergic, endocannabinoid and plasticity-related genes in the dorsomedial prefrontal cortex after sucrose self-administration, which was more pronounced after 30 days of withdrawal (Roura-Martínez et al., 2019). The relationship of the PNN modulations here reported with the aforementioned changes in neurotransmitter receptors is unclear at the moment. However, interactions between AMPA receptor function and PNN levels have been reported (Chen et al., 2016) which could be especially relevant in the case of sucrose seeking incubation where decrements in the GluA1 subunit of the AMPA glutamate receptors are observed (Roura-Martínez et al., 2019).

### Opposite changes in perineuronal net content in the insula

The insula appears to be involved in reinforcer seeking after self-administration protocols that induce incubation. For example, it has been shown to mediate methamphetamine seeking, at least after protracted withdrawal (Li et al., 2018; Venniro et al., 2017) and, with regard to alcohol, Abdolahi et al., (2010) observed interesting modulations throughout the withdrawal in the pattern of phosphorylation of the dopamine-related protein DARPP-34. In this work, we have found opposite, and hemisphere-specific alterations in the PNN content in the insula after the self-administration of heroin (a decrease in the right hemisphere), and sucrose (an increase in the left hemisphere) that were evident even after 30 days of withdrawal. Coincidentally with this later effect, Chen et al., (2015) showed that PNNs increase in the left granular insula after alcohol self-administration although this modulation was only evident at early withdrawal. It is now necessary to elucidate if the insular PNNs that decrease after heroin self-administration share a similar functional profile with those that increase after sucrose self-administration and also to gain a deeper understanding of the importance of the hemisphere specificity in these functions, an issue that remains completely unexplored.

### Lack of changes in perineuronal net content in OFC

We have not found any change in the PNN content either in the ventral or in the lateral OFCs. This lack of effects is supported in part by the literature. Indeed, while a high-fat diet (Dingess et al., 2018) or an intragastric administration of alcohol (Coleman et al., 2014) modified PNN content in the OFC, previous studies that employed self-administration and withdrawal paradigms, similar to the ones used here, showed no changes with nicotine (Vazquez-Sanroman et al., 2017) or sucrose (Slaker et al., 2016).

### Concluding remarks

The problem of relapse in addictive disorders remains unsolved. This is due, in part, to the lack of a complete understanding of the general mechanisms that govern relapse across the different types of addictions, which likely involve learning and memory processes. PNNs are important structures that regulate synaptic plasticity and neuronal function and as such, are potential actors in relapse. Here, we have provided a much-needed comparative analysis of PNN dynamics in a self-administration model that allows the examination of the neurobiological alterations across the time course of abstinence. The down-regulation of PNNs appeared as a common effect during abstinence but, in certain regions, there were drug-specific effects that call for a more detailed analysis. We suggest that, if we are to fully understand the intricate network of mechanisms that operate in the relapse phenomenon, this sort of comparative analysis using self-administration models with different drugs and natural rewards, should be widely adopted in the future.

## Acknowledgements

We would like to thank Rosa Ferrado, Alberto Marcos and Gonzalo Moreno for excellent technical assistance and Lidia Blázquez-Llorca for helping to set up the histochemical procedures used in this work. The Department of Psychobiology of UNED has provided continuous support. The present work has been funded by the Spanish Ministry of Economy and Competitiveness (PSI2016-80541-P to Emilio Ambrosio and Alejandro Higuera-Matas); Spanish Ministry of Health, Social Services and Equality (Network of Addictive Disorders – Project n°: RTA-RD16/0017/0022 of the Institute of Health Carlos III to Emilio Ambrosio and Plan Nacional Sobre Drogas, Project n°: 2016I073 to Emilio Ambrosio and 2017I042 to Alejandro Higuera-Matas);UNED (Plan for the Promotion of Research to Emilio Ambrosio and Alejandro Higuera-Matas); the European Union (Project n°: JUST/2017/AG-DRUG-806996-JUSTSO to Emilio Ambrosio) and the BBVA Foundation (2017 Leonardo Grant for Researchers and Cultural Creators to Alejandro Higuera Matas). Also, part of this work was supported by the Cajal Blue Brain Platform (CSIC/UPM, Spain). Marcos Ucha received a predoctoral fellowship awarded by the Ministry of Science and Innovation BES-2011-043814) and David Roura-Martínez received a predoctoral fellowship from the Plan for the Promotion of Research of UNED.

## REFERENCES

Abdolahi, A., Acosta, G., Breslin, F.J., Hemby, S.E., Lynch, W.J., 2010. Incubation of nicotine seeking is associated with enhanced protein kinase A-regulated signaling of dopamine- and cAMP-regulated phosphoprotein of 32 kDa in the insular cortex. Eur. J. Neurosci. https://doi.org/10.1111/j.1460-9568.2010.07114.x

Aoyama, K., Barnes, J., Grimm, J.W., 2014. Incubation of saccharin craving and within-session changes in responding for a cue previously associated with saccharin. Appetite. https://doi.org/10.1016/j.appet.2013.10.003

Bedi, G., Preston, K.L., Epstein, D.H., Heishman, S.J., Marrone, G.F., Shaham, Y., de Wit, H., 2011. Incubation of cue-induced cigarette craving during abstinence in human smokers. Biol. Psychiatry 69, 708–11. https://doi.org/10.1016/j.biopsych.2010.07.014

Ben-Shahar, O., Obara, I., Ary, A.W., Ma, N., Mangiardi, M.A., Medina, R.L., Szumlinski, K.K., 2009. Extended daily access to cocaine results in distinct alterations in Homer 1b/c and NMDA receptor subunit expression within the medial prefrontal cortex. Synapse 63, 598–609. https://doi.org/10.1002/syn.20640

Ben-Shahar, O., Sacramento, A.D., Miller, B.W., Webb, S.M., Wroten, M.G., Silva, H.E., Caruana, A.L., Gordon, E.J., Ploense, K.L., Ditzhazy, J., Kippin, T.E., Szumlinski, K.K., 2013. Deficits in Ventromedial Prefrontal Cortex Group 1 Metabotropic Glutamate Receptor Function Mediate Resistance to Extinction during Protracted Withdrawal from an Extensive History of Cocaine Self-Administration. J. Neurosci. https://doi.org/10.1523/JNEUROSCI.3710-12.2013

Bienkowski, P., Rogowski, A., Korkosz, A., Mierzejewski, P., Radwanska, K., Kaczmarek, L., Bogucka-Bonikowska, A., Kostowski, W., 2004. Time-dependent changes in alcohol-seeking behaviour during abstinence. Eur. Neuropsychopharmacol. https://doi.org/10.1016/j.euroneuro.2003.10.005

Blackwood, C.A., Hoerle, R., Leary, M., Schroeder, J., Job, M.O., McCoy, M.T., Ladenheim, B., Jayanthi, S., Cadet, J.L., 2018. Molecular Adaptations in the Rat Dorsal Striatum and Hippocampus Following Abstinence-Induced Incubation of Drug Seeking After Escalated Oxycodone Self-Administration. Mol. Neurobiol. https://doi.org/10.1007/s12035-018-1318-z

Bossert, J.M., Ghitza, U.E., Lu, L., Epstein, D.H., Shaham, Y., 2005. Neurobiology of relapse to heroin and cocaine seeking: An update and clinical implications, in: European Journal of Pharmacology. Elsevier, pp. 36–50. https://doi.org/10.1016/j.ejphar.2005.09.030

Chen, H., He, D., Lasek, A.W., 2015. Repeated Binge Drinking Increases Perineuronal Nets in the Insular Cortex. Alcohol. Clin. Exp. Res. https://doi.org/10.1111/acer.12847

Chen, W., Li, Y.S., Gao, J., Lin, X.Y., Li, X.H., 2016. AMPA receptor antagonist NBQX decreased seizures by normalization of perineuronal nets. PLoS One 11, e0166672. https://doi.org/10.1371/journal.pone.0166672

Coleman, L.G., Liu, W., Oguz, I., Styner, M., Crews, F.T., 2014. Adolescent binge ethanol treatment alters adult brain regional volumes, cortical extracellular matrix protein and behavioral flexibility. Pharmacol. Biochem. Behav. https://doi.org/10.1016/j.pbb.2013.11.021

Dingess, P.M., Harkness, J.H., Slaker, M., Zhang, Z., Wulff, S.S., Sorg, B.A., Brown, T.E., 2018. Consumption of a High-Fat Diet Alters Perineuronal Nets in the Prefrontal Cortex. Neural Plast. https://doi.org/10.1155/2018/2108373

Everitt, B.J., Robbins, T.W., 2016. Drug Addiction: Updating Actions to Habits to Compulsions Ten Years On. Annu. Rev. Psychol. 67, 150807174122003. https://doi.org/10.1146/annurev-psych-122414-033457

George, O., Koob, G.F., 2010. Individual differences in prefrontal cortex function and the transition from drug use to drug dependence. Neurosci. Biobehav. Rev. 35, 232–47. https://doi.org/10.1016/j.neubiorev.2010.05.002

Gould, A.T., Sacramento, A.D., Wroten, M.G., Miller, B.W., Von Jonquieres, G., Klugmann, M., Ben-Shahar, O., Szumlinski, K.K., 2015. Cocaine-elicited imbalances in ventromedial prefrontal cortex Homer1 versus Homer2 expression: Implications for relapse. Addict. Biol. https://doi.org/10.1111/adb.12088

Grimm, J.W., Hope, B.T., Wise, R.A., Shaham, Y., 2001. Incubation of cocaine craving after withdrawal. Nature 412, 141–142. https://doi.org/10.1038/35084134

Grimm, J.W., Shaham, Y., Hope, B.T., 2002. Effect of cocaine and sucrose withdrawal period on extinction behavior, cue-induced reinstatement, and protein levels of the dopamine transporter and tyrosine hydroxylase in limbic and cortical areas in rats. Behav. Pharmacol. 13, 379–388.

Kirschmann, E.K., Pollock, M.W., Nagarajan, V., Torregrossa, M.M., 2017. Effects of Adolescent Cannabinoid Self-Administration in Rats on Addiction-Related Behaviors and Working Memory. Neuropsychopharmacology 42, 989–1000. https://doi.org/10.1038/npp.2016.178

Koya, E., Uejima, J.L., Wihbey, K.A., Bossert, J.M., Hope, B.T., Shaham, Y., 2009. Role of ventral medial prefrontal cortex in incubation of cocaine craving. Neuropharmacology 56, 177–185. https://doi.org/10.1016/j.neuropharm.2008.04.022

Krasnova, I.N., Marchant, N.J., Ladenheim, B., McCoy, M.T., Panlilio, L. V., Bossert, J.M., Shaham, Y., Cadet, J.L., 2014. Incubation of methamphetamine and palatable food craving after punishment-induced abstinence. Neuropsychopharmacology 39, 2008–2016. https://doi.org/10.1038/npp.2014.50

Lasek, A.W., Chen, H., Chen, W.Y., 2018. Releasing Addiction Memories Trapped in Perineuronal Nets. Trends Genet. https://doi.org/10.1016/j.tig.2017.12.004

Li, P., Wu, P., Xin, X., Fan, Y.L., Wang, G. Bin, Wang, F., Ma, M.Y., Xue, M.M., Luo, Y.X., Yang, F. De, Bao, Y.P., Shi, J., Sun, H.Q., Lu, L., 2015. Incubation of alcohol craving during abstinence in patients with alcohol dependence. Addict. Biol. https://doi.org/10.1111/adb.12140

Li, X., Witonsky, K., Lofaro, O.M., Surjono, F., Zhang, J., Bossert, J.M., Shaham, Y., 2018. Role of anterior intralaminar nuclei of thalamus projections to dorsomedial striatum in incubation of methamphetamine craving. J. Neurosci. https://doi.org/10.1523/JNEUROSCI.2873-17.2018

Li, X., Zeric, T., Kambhampati, S., Bossert, J.M., Shaham, Y., 2015. The Central Amygdala Nucleus is Critical for Incubation of Methamphetamine Craving. Neuropsychopharmacology 40, 1297–1306. https://doi.org/10.1038/npp.2014.320

Miller, B.W., Wroten, M.G., Sacramento, A.D., Silva, H.E., Shin, C.B., Vieira, P.A., Ben-Shahar, O., Kippin, T.E., Szumlinski, K.K., 2017. Cocaine craving during protracted withdrawal requires PKCε priming within vmPFC. Addict. Biol. https://doi.org/10.1111/adb.12354

Mizoguchi, H., Yamada, K., Mouri, A., Niwa, M., Mizuno, T., Noda, Y., Nitta, A., Itohara, S., Banno, Y., Nabeshima, T., 2007. Role of matrix metalloproteinase and tissue inhibitor of MMP in methamphetamine-induced behavioral sensitization and reward: Implications for dopamine receptor down-regulation and dopamine release. J. Neurochem. https://doi.org/10.1111/j.1471-4159.2007.04623.x

Müller Ewald, V.A., De Corte, B.J., Gupta, S.C., Lillis, K. V., Narayanan, N.S., Wemmie, J.A., LaLumiere, R.T., 2018. Attenuation of cocaine seeking in rats via enhancement of infralimbic cortical activity using stable step-function opsins. Psychopharmacology (Berl). https://doi.org/10.1007/s00213-018-4964-y

O’Brien, C.P., 2008. Review. Evidence-based treatments of addiction. Philos. Trans. R. Soc. Lond. B. Biol. Sci. 363, 3277–86. https://doi.org/10.1098/rstb.2008.0105

Parvaz, M.A., Moeller, S.J., Goldstein, R.Z., 2016. Incubation of cue-induced craving in adults addicted to cocaine measured by electroencephalography. JAMA Psychiatry. https://doi.org/10.1001/jamapsychiatry.2016.2181

Reichelt, A.C., Gibson, G.D., Abbott, K.N., Hare, D.J., 2019. A high-fat high-sugar diet in adolescent rats impairs social memory and alters chemical markers characteristic of atypical neuroplasticity and parvalbumin interneuron depletion in the medial prefrontal cortex. Food Funct. https://doi.org/10.1039/c8fo02118j

Roura-Martínez, D., Ucha, M., Orihuel, J., Castillo, C.A., Marcos, A., Ambrosio, E., Matas, A.H., 2019. Central nucleus of the amygdala as a common substrate of the incubation of drug and natural reinforcer seeking. Addict. Biol. 1–12. https://doi.org/10.1111/adb.12706

Shalev, U., Morales, M., Hope, B., Yap, J., Shaham, Y., 2001. Time-dependent changes in extinction behavior and stress-induced reinstatement of drug seeking following withdrawal from heroin in rats. Psychopharmacology (Berl). https://doi.org/10.1007/s002130100748

Shepard, J.D., Bossert, J.M., Liu, S.Y., Shaham, Y., 2004. The anxiogenic drug yohimbine reinstates methamphetamine seeking in a rat model of drug relapse. Biol. Psychiatry. https://doi.org/10.1016/j.biopsych.2004.02.032

Shin, C.B., Templeton, T.J., Chiu, A.S., Kim, J., Gable, E.S., Vieira, P.A., Kippin, T.E., Szumlinski, K.K., 2018. Endogenous glutamate within the prelimbic and infralimbic cortices regulates the incubation of cocaine-seeking in rats. Neuropharmacology 128, 293–300. https://doi.org/10.1016/j.neuropharm.2017.10.024

Slaker, M., Barnes, J., Sorg, B.A., Grimm, J.W., 2016. Impact of environmental enrichment on perineuronal nets in the prefrontal cortex following early and late abstinence from sucrose self-administration in rats. PLoS One 11. https://doi.org/10.1371/journal.pone.0168256

Slaker, M., Churchill, L., Todd, R.P., Blacktop, J.M., Zuloaga, D.G., Raber, J., Darling, R.A., Brown, T.E., Sorg, B.A., 2015. Removal of perineuronal nets in the medial prefrontal cortex impairs the acquisition and reconsolidation of a cocaine-induced conditioned place preference memory. J. Neurosci. https://doi.org/10.1523/JNEUROSCI.3592-14.2015

Sorg, B.A., Berretta, S., Blacktop, J.M., Fawcett, J.W., Kitagawa, H., Kwok, J.C.F., Miquel, M., 2016. Casting a wide net: Role of perineuronal nets in neural plasticity, in: Journal of Neuroscience. https://doi.org/10.1523/JNEUROSCI.2351-16.2016

Swinford-Jackson, S.E., Anastasio, N.C., Fox, R.G., Stutz, S.J., Cunningham, K.A., 2016. Incubation of cocaine cue reactivity associates with neuroadaptations in the cortical serotonin (5-HT) 5-HT2Creceptor (5-HT2CR) system. Neuroscience 2, 50–61. https://doi.org/10.1016/j.neuroscience.2016.02.052

Szumlinski, K.K., Ary, A.W., Shin, C.B., Wroten, M.G., Courson, J., Miller, B.W., Ruppert-Majer, M., Hiller, J.W., Shahin, J.R., Ben-Shahar, O., Kippin, T.E., 2018. PI3K activation within ventromedial prefrontal cortex regulates the expression of drug-seeking in two rodent species. Addict. Biol. https://doi.org/10.1111/adb.12696

Van den Oever, M.C., Lubbers, B.R., Goriounova, N. a, Li, K.W., Van der Schors, R.C., Loos, M., Riga, D., Wiskerke, J., Binnekade, R., Stegeman, M., Schoffelmeer, A.N.M., Mansvelder, H.D., Smit, A.B., De Vries, T.J., Spijker, S., 2010. Extracellular matrix plasticity and GABAergic inhibition of prefrontal cortex pyramidal cells facilitates relapse to heroin seeking. Neuropsychopharmacology 35, 2120–33. https://doi.org/10.1038/npp.2010.90

Vazquez-Sanroman, D.B., Monje, R.D., Bardo, M.T., 2017. Nicotine self-administration remodels perineuronal nets in ventral tegmental area and orbitofrontal cortex in adult male rats. Addict. Biol. https://doi.org/10.1111/adb.12437

Venniro, M., Caprioli, D., Zhang, M., Whitaker, L.R., Zhang, S., Warren, B.L., Cifani, C., Marchant, N.J., Yizhar, O., Bossert, J.M., Chiamulera, C., Morales, M., Shaham, Y., 2017. The Anterior Insular Cortex→Central Amygdala Glutamatergic Pathway Is Critical to Relapse after Contingency Management. Neuron 96, 414–427. https://doi.org/10.1016/j.neuron.2017.09.024

Wang, G., Shi, J., Chen, N., Xu, L., Li, J., Li, P., Sun, Y., Lu, L., 2013. Effects of length of abstinence on decision-making and craving in methamphetamine abusers. PLoS One 8, e68791. https://doi.org/10.1371/journal.pone.0068791

